## Supplemental Figures for "Text mining-based word representations for biomedical data analysis and machine learning tasks"

**Supplemental Figure 1**

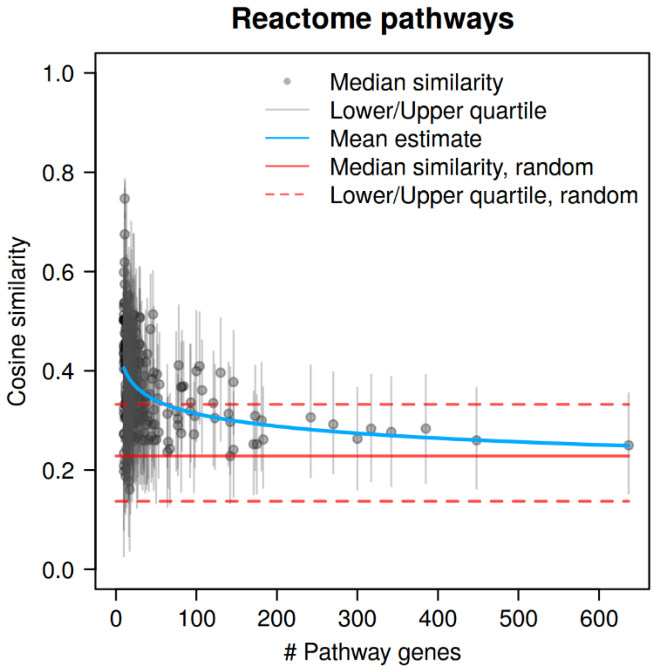


**S1 Fig. Gene-gene cosine similarities within Reactome pathways of given number of genes.** Median, lower and upper quartiles are presented for a random sample of 2000 gene pairs. A mean trend was estimated using the function f(x) = (xa + b)-1.

**Supplemental Figure 2**

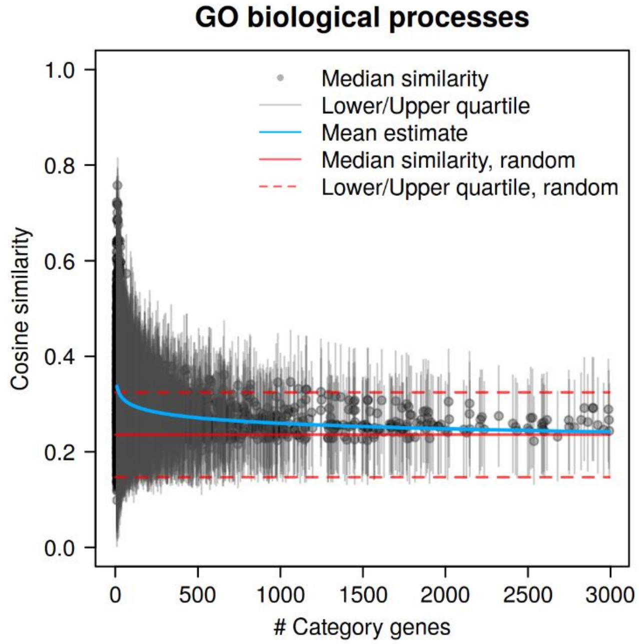


**S2 Fig. Gene-gene cosine similarities within GO biological processes of given number of genes.** Median, lower and upper quartiles are presented for a random sample of 2000 gene pairs. A mean trend was estimated using the function f(x) = (xa + b)-1.

**Supplemental Figure 3**

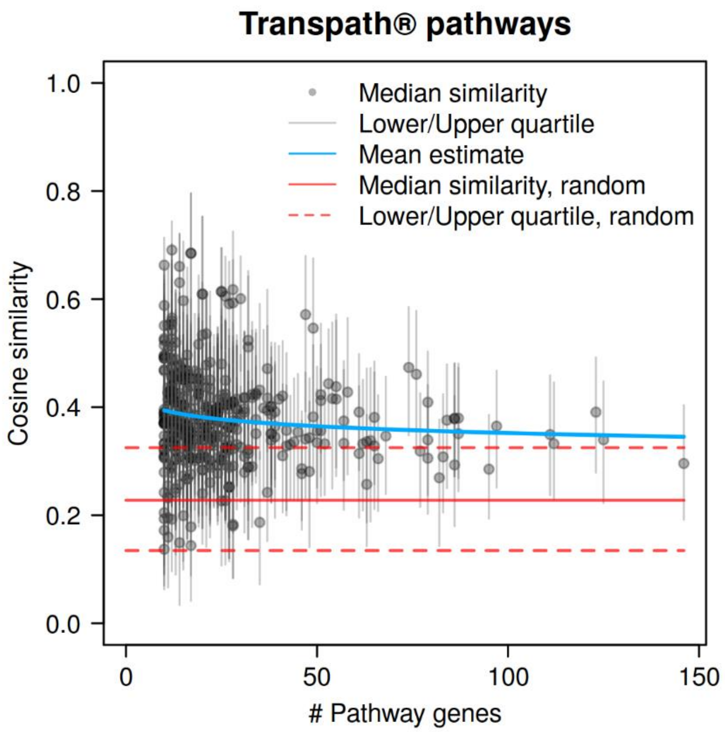


**S3 Fig. Gene-gene cosine similarities within TRANSPATH® pathways of given number of genes.** Median, lower and upper quartiles are presented for a random sample of 2000 gene pairs. A mean trend was estimated using the function f(x) = (xa + b)-1.

**Supplemental Figure 4**

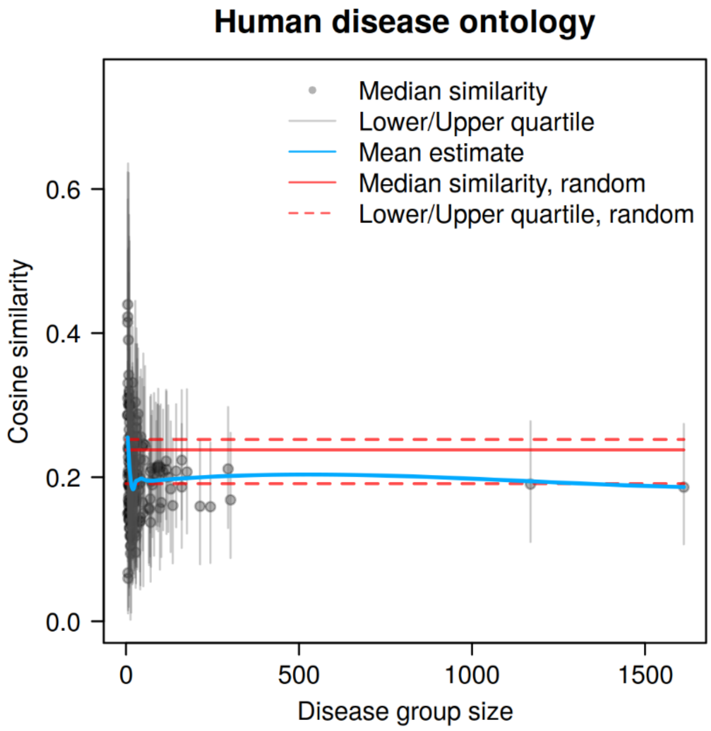


**S4 Fig. Disease-disease cosine similarities within human disease ontology groups of given number of diseases.** Median, lower and upper quartiles are presented for a random sample of 700 disease pairs. A mean trend was estimated by non-parametric local regression (Loess).

**Supplemental Figure 5**

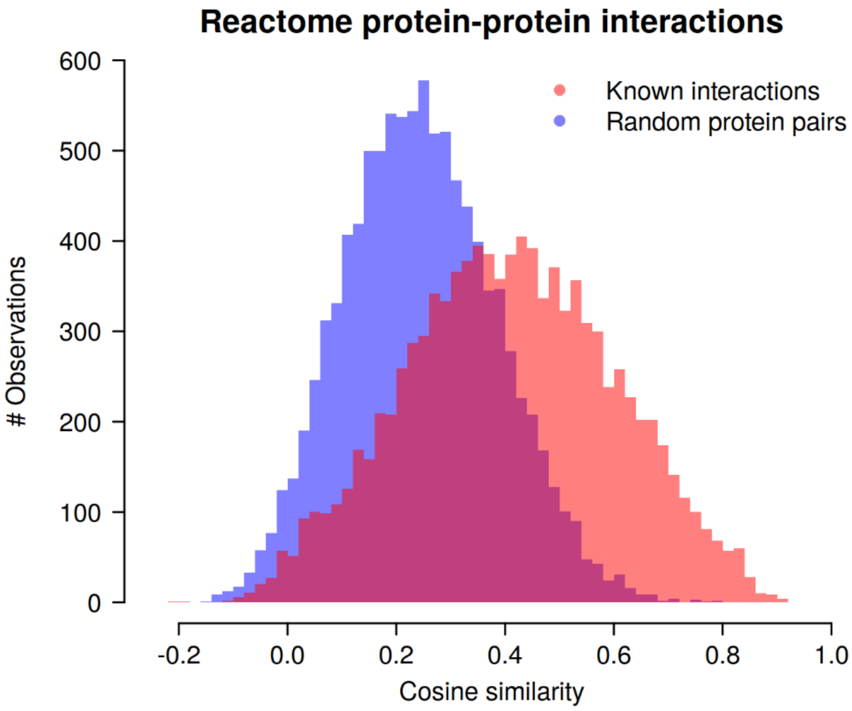


**S5 Fig. Histograms of genes with or without known Reactome protein-protein interactions.** Each group contained a random sample of 10000 pairs.

**Supplemental Figure 6**

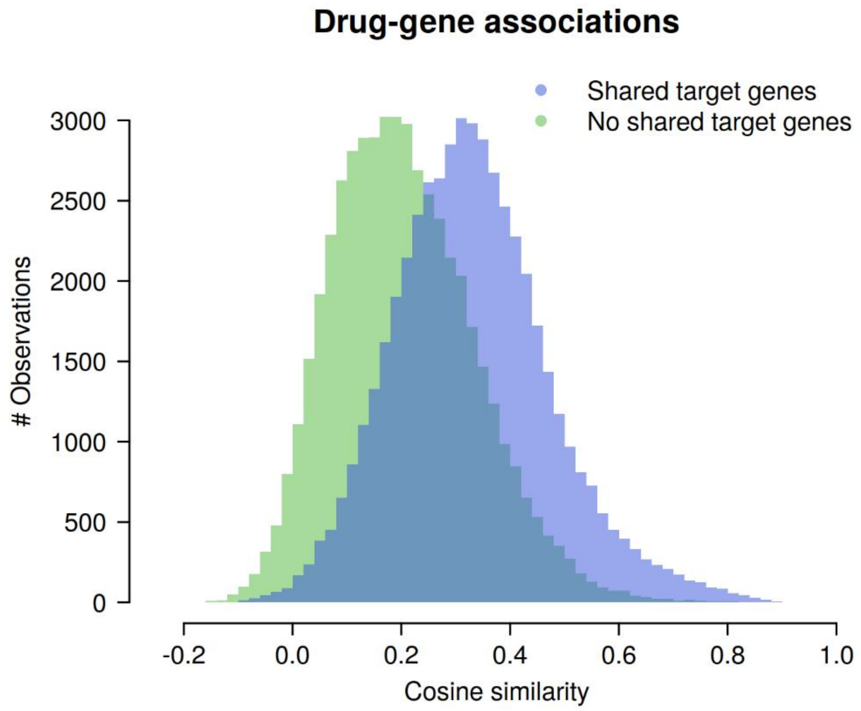


**S6 Fig. Histograms of drug-gene associations with or without shared target genes.** Each group contained a random sample of 50000 drug pairs.

**Supplemental Figure 7**

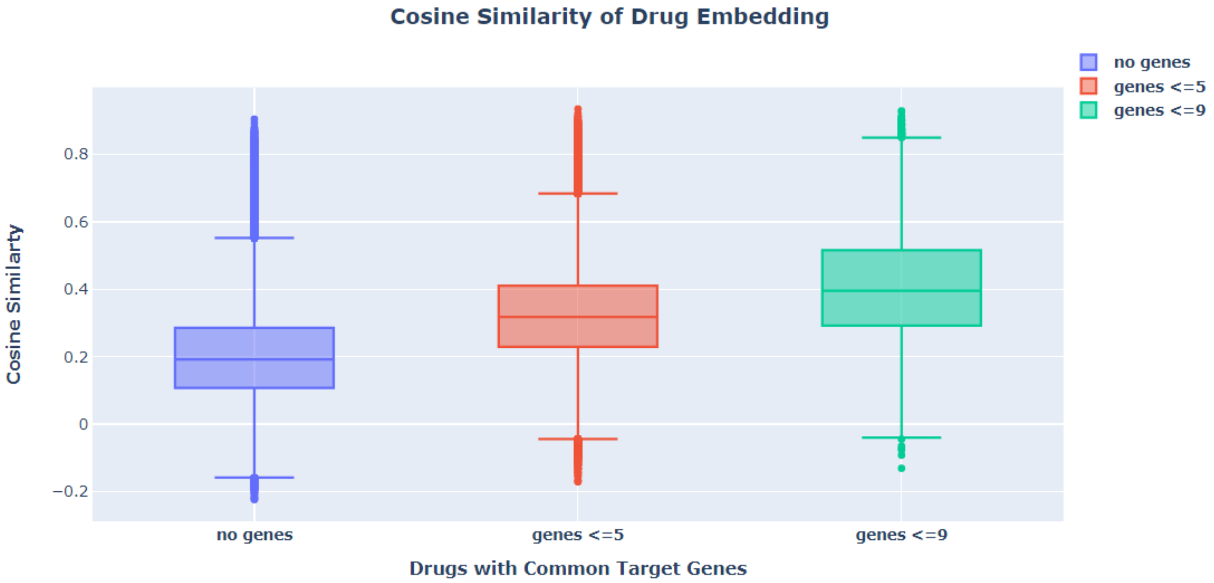


**S7 Fig. Drug-drug cosine similarity distributions with shared genes of given number in DrugBank.** Drug-drug groups were estimated by counting the number of shared genes between two drugs presented in the embedding. Group1 (no genes: median= 0.192, lower quartile = 0.108, upper quartile = 0,286), group2 (genes ≤ 5: median= 0.318, lower quartile = 0.229, upper quartile = 0,411), group3 (genes ≤ 9: median= 0.396, lower quartile = 0.292, upper quartile = 0,516).
